# The *In Situ* Structure and Distribution of V-ATPase in *C. elegans* Apical Membrane Stacks

**DOI:** 10.64898/2025.12.19.695382

**Authors:** Euan Pyle, Anna M. Steyer, Anastasiia Babenko, Higor Vinícius Dias Rosa, Simone Mattei

**Author notes:** Correspondence should be addressed to Simone Mattei.

## Abstract

The cuticle of *Caenorhabditis elegans* is an apical extracellular matrix composed primarily of collagens which protects the organism from external stresses and facilitates mobility. The cuticle directly contacts several epithelial cell structures including regular, repeating folds of the apical plasma membrane, known as apical membrane stacks (AMSs). Here, we examine the structure of AMSs using *in situ* cryo-electron tomography (cryo-ET). We find that V-ATPases are distributed across the cytoplasmic-facing membranes of the AMSs. We then determine the *in situ* structure of V-ATPase and show that it is highly conserved relative to other eukaryotic V-ATPases. Analysis of the distribution of V-ATPases on the AMSs shows that they form clusters in which individual V-ATPases are tightly packed but do not oligomerise. Our data also show that complete, fully assembled V-ATPases line the AMSs. Previous studies have established that V-ATPases can only act as ATP-dependent proton pumps in their fully assembled state. Our findings indicate that V-ATPases arranged along the AMSs pump protons directly into the extracellular matrix, suggesting that a primary function of AMSs is to acidify the cuticle.

## Introduction

The cuticle in *Caenorhabditis elegans* is an apical extracellular matrix that covers the entire surface of the organism and acts as a barrier protecting it from external biotic and abiotic stresses. Additionally, the cuticle maintains the body shape of the organism and assists mobility through its role as an external skeleton^1^. The cuticle is composed primarily of collagens, which form insoluble and highly ordered complexes^1,2^. It is produced by epithelial cells and is attached to the basal lamina and hypodermis through hemidesmosomes^1^. At each larval stage of the *C. elegans* lifecycle, a new cuticle is synthesised while the old cuticle is broken down and shed to allow growth^1^. Many genes have been identified that play a role in cuticle and collagen degradation^3^, but the precise molecular details of this process remain unclear.

Previous studies have described the presence of regular, repeating folds in the apical plasma membrane of the epidermis in *C. elegans*^4–8^. These structures are known as either apical membrane stacks (AMSs)^5,7^ or meisosomes^8^. Analysis of room-temperature transmission electron microscopy (TEM) data has described and characterised their organisation and ultrastructure^8^. Generally, AMSs consist of up to 30 membrane folds^8^, resulting in rows of parallel membranes that form a large interface with the cuticle. The gaps between the parallel membranes are regularly spaced, with distances of 35 nm separating cytoplasm-facing membranes and 20 nm separating cuticle-facing membranes. They are often found in close proximity to both mitochondria^8^ and multivesicular bodies^7^ (MVBs). AMSs are primarily found in the epidermal syncytium hyp7 at the lateral, dorsal, and ventral ridges in all larval stages and adults.

Whilst the general ultrastructure and localisation of the AMSs is well characterised, their function and molecular composition remain debated. Previous studies have established that VHA-5, a subunit of the transmembrane domain of the vacuolar ATPase (V-ATPase), and RAL-1, a GTPase, both co-localise with AMSs^5,7^. Whilst both VHA-5 and RAL-1 have been implicated in the process of exosome secretion into the cuticle^5^, the functional roles of these proteins in AMSs have not been fully established.

*C. elegans* has recently emerged as a model system for *in situ* structural biology of multicellular organisms by cryo-electron tomography (cryo-ET)^9–11^. In this study, we used cryo-ET to image and analyse *C. elegans in situ*. Our high-resolution cryo-ET data allowed us to visualise native AMSs which showed distinct protein molecules lining their surface. Subtomogram averaging revealed that these particles were V-ATPases. We then analysed the structure and distribution of the V-ATPases within the AMSs, which suggested that cuticle acidification is a possible functional role for AMSs in *C. elegans*.

## Results

### Cryo-electron tomography visualises AMSs

To visualise *C. elegans in situ*, we carried out serial cryo-lift out^10^ on a high-pressure frozen L4-stage worm before imaging by cryo-ET (Figure 1). We utilised the fluorescence from mMaple-tagged chitin synthase-2 (CHS-2), which is known to localise to the pharynx^12^, to target this region during the lift out process (Figure 1a-d). After lift out, we carried out fluorescence microscopy on the resulting serial slices of *C. elegans*, known as lamellae. Fluorescence data confirmed the presence of pharynx-localised CHS-2 in the majority of our lamellae (Supplementary Figure 1). Interestingly, CHS-2 was expressed only in the pharynx muscle cells and not in the pharynx marginal cells (Supplementary Figure 1).

**Figure 1:**
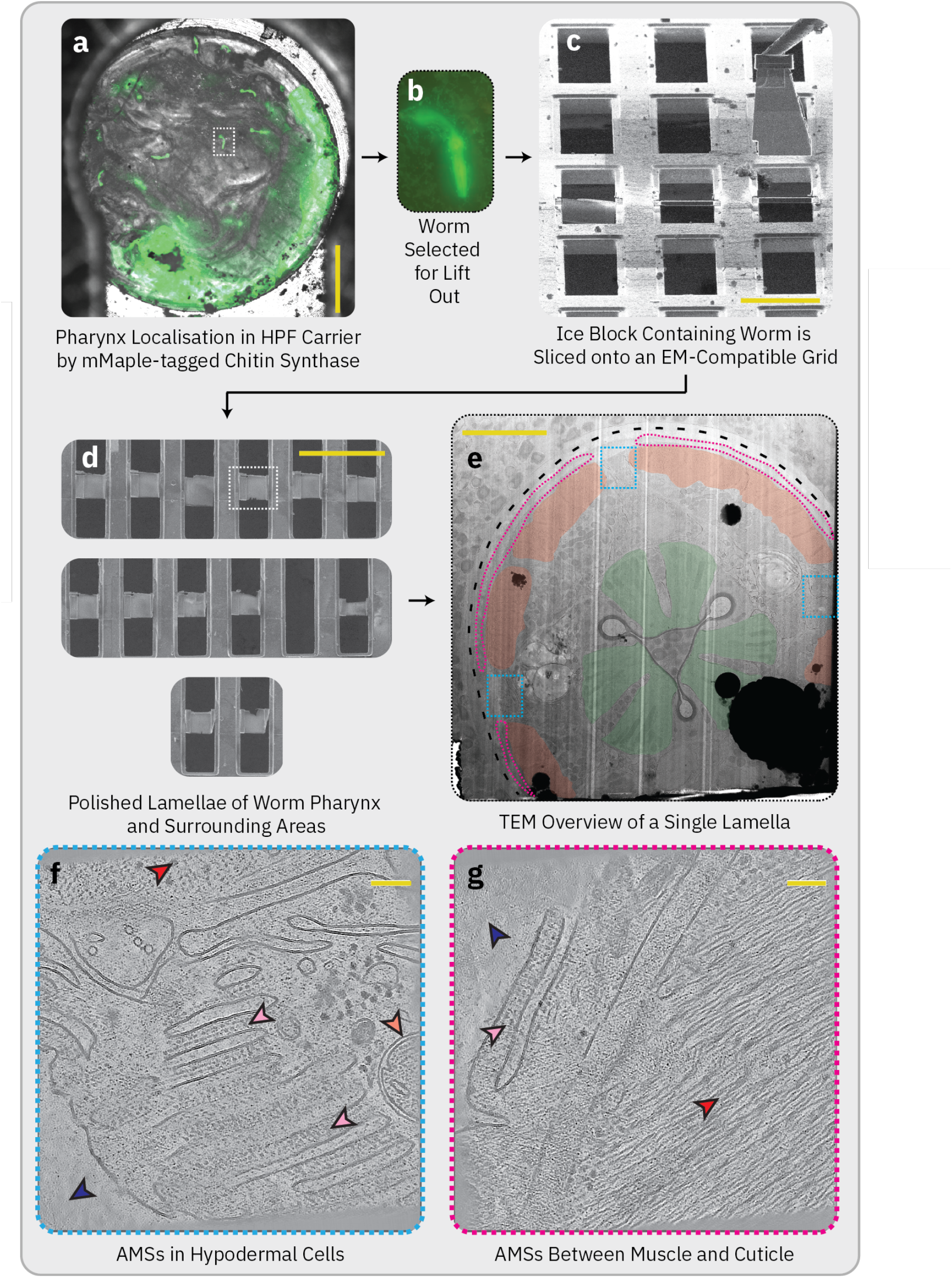
Workflow for serial lift out of *C. elegans*, TEM of one of the produced lamellae, and tomograms of regions containing AMSs. **(a)** Widefield microscopy image of a high-pressure frozen carrier containing *C. elegans* with overlaid cryo-fluorescence signal of CHS-2::mMaple to localise the pharynx of the worms. A white box indicates the worm selected to be lifted out of the sample and imaged using cryo-ET. Scale bar = 500 μm. **(b)** Inset of the fluorescence signal of the worm selected for lift out **(c)** FIB image of the lifted-out block, containing the worm visualised in *b*, above the recipient grid on which the lamellae were deposited. A successfully deposited lamella (pre-thinning) can be seen in the middle of the image. Scale bar = 50 μm. **(d)** SEM image of the lamellae deposited on the recipient grid after thinning the ice to a target thickness of 200 nm. A white box indicates the lamella shown in *e*. Scale bar = 100 μm. **(e)** TEM overview of the lamella indicated in *d*. The dotted black line indicates the cuticle, the regions highlighted in red correspond to muscle cells, the regions highlighted in green correspond to the pharynx muscle cells, the dotted blue boxes indicate the regions in hypodermal cells in which the majority of AMSs were found, and the regions encircled by the dotted pink lines correspond to the region in which AMSs were found between muscle cells and cuticle. Scale bar = 5 μm. **(f)** A tomogram of AMSs found in hypodermal cells. The pink arrows indicate AMSs, the red arrow indicates a neighbouring muscle cell, the blue arrow indicates the cuticle, and the orange arrow indicates a nearby mitochondrion. Scale bar = 100 nm. **(g)** A tomogram of AMSs found in the region between muscle cells and cuticle. Arrows indicate the same objects as in *f*. Scale bar = 100 nm.

In total, we generated 11 lamellae of the pharynx and its surrounding regions that were of sufficient quality for imaging by cryo-ET. From these lamellae, we collected a total of 216 tomograms. Within 8 of these tomograms, from 7 different lamellae, we found clear evidence of plasma membrane folds next to the cuticle which visually strongly resembled AMSs^8^ (Figure 1f-g; Supplementary Figure 2). Segmentation analysis of these folds revealed that they have similar approximate dimensions as AMSs, with regularly spaced gaps of 44 nm (SD = 0.88) separating cytoplasmic-facing membranes, and 31 nm (SD = 1.93) separating cuticle-facing membranes^8^ (Supplementary Figure 3). Additionally, we often found mitochondria in the immediate vicinity of the folds, as previously reported^8^ (Figure 1f). Together, these data demonstrate that the observed plasma membrane folds are AMSs. AMSs were almost entirely found within hypodermal cells in the dorsal, ventral, and/or lateral ridge regions (Figure 1f). We also found one instance of an AMS located at the plasma membrane in a hypodermal cell wedged between muscle and the cuticle (Figure 1g).

### AMSs within their cellular context

The function of the AMSs in *C. elegans* is unclear. To understand their role, we analysed the position of the AMSs within their cellular context by tomogram segmentation (Figure 2). Segmentation confirmed previously reported data that AMSs are formed by folds in the plasma membrane and that mitochondria are often closely associated with them^8^. In addition to the mitochondria, we also noted the proximity of collagen in the cuticle to the AMSs (Figure 2a). The collagen appears in its ordered form, as shown by the characteristic V-shaped striations of collagen in the basal layer^13,14^ (Figure 2a). Collagen closer to the AMSs appears disordered; however, this apparent loss of order may result from collagen fibrils being oriented along the Z axis of the tomogram, which obscures their arrangement.

**Figure 2:**
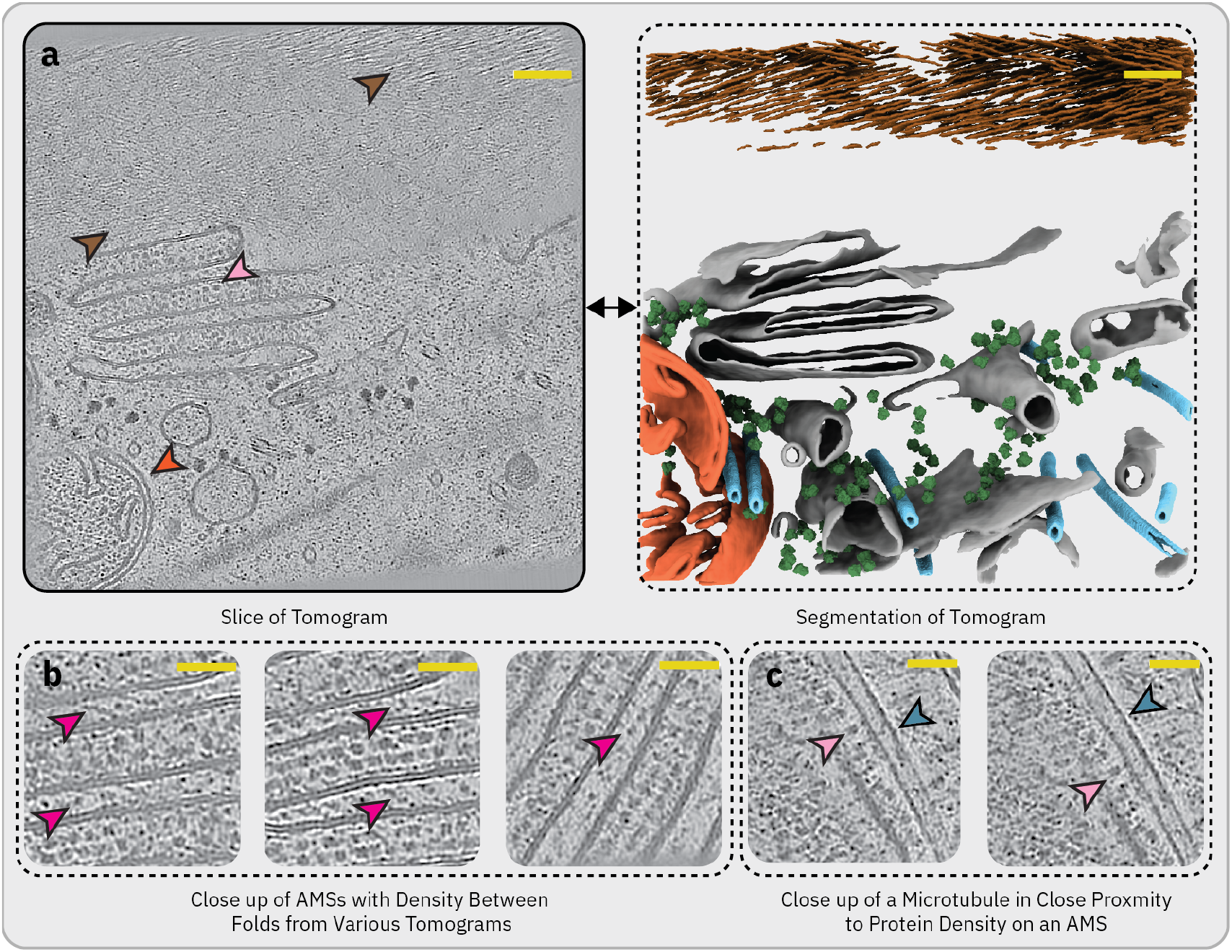
AMSs within their cellular context. **(a)** Tomogram containing AMSs (left) where the light pink arrow indicates AMSs, the brown arrow indicates collagen in the cuticle, and the orange arrow indicates a nearby mitochondrion. The resulting segmentation of this tomogram (right) displays ordered collagen in brown, mitochondrial membrane in orange, membranes in grey, microtubules in light blue, and ribosomes in green. Note the plasma membrane is incomplete due to the missing wedge from cryo-ET data collection obscuring views of the membrane bilayer. Scale bar = 100 nm. **(b)** Close up images from various tomograms with purple arrows indicating unidentified densities between AMSs. Scale bar = 50 nm. **(c)** Close up images from a tomogram with pink arrows indicating protein density on the AMSs, and blue arrows indicating a microtubule in close proximity. Scale bar = 50 nm.

We consistently observed many visually identical, membrane-bound protein densities on the cytoplasmic side of the AMSs (Figure 1f-g; Figure 2a). These protein densities were previously reported^8^ but were not visualised at a resolution sufficient to distinguish individual particles and, consequently, had not been identified. Additionally, we consistently observed density between the AMSs on the extracellular side of the membrane (Figure 2b). The signal-to-noise ratio of the tomograms, and the small size of the densities, made these objects challenging to quantify or generate a subtomogram average of. However, we did note multiple instances of the densities stretching from one fold of the membrane stack to another (Figure 2b). We also observed a microtubule in close proximity to the protein molecules lining the AMSs in one tomogram with density seemingly connecting the microtubule and the membrane protein (Figure 2c).

### The *in situ* map of V-ATPase in AMSs

We carried out a subtomogram averaging workflow using Warp/M^15,16^ and RELION^17,18^ to obtain a reconstruction of the particles lining the cytoplasmic-facing membranes of the AMSs. Initially, we generated a subtomogram average of the *C. elegans* ribosome from tomograms both with and without AMSs to 5.6 Å resolution (Supplementary Figure 4). We used this map to improve the quality of the tomograms by refining tilt-series alignment in M^16^ using the ribosomes as quasi-fiducial markers (see Materials and Methods). After the improved tomograms were generated, we produced a subtomogram average of the particles lining the AMSs. The resulting 17.4 Å map was unambiguously the *C. elegans* V-ATPase (Figure 3).

**Figure 3:**
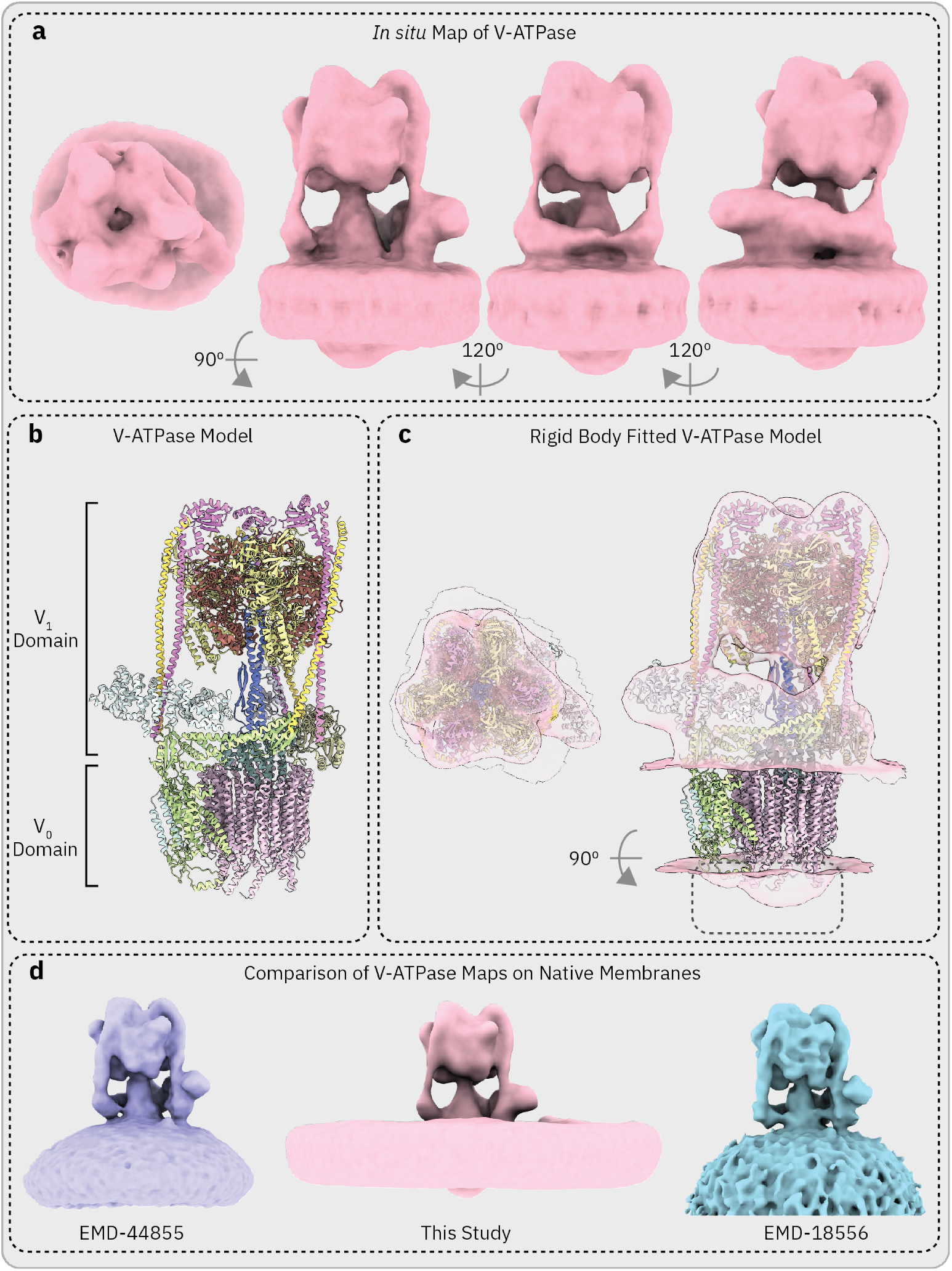
Structure of the V-ATPase from *C. elegans* AMSs. **(a)** 17.4 Å density map of V-ATPase obtained *in situ* from *C. elegans* AMSs viewed from four different angles. **(b)** Model of *C. elegans* V-ATPase. The structures of individual subunits were obtained by AlphaFold predictions, which were superimposed on a pre-existing V-ATPase structure (PDB: 9BRA) before fitting into our density map. VHA-19 (Ac45) could not be accurately fitted into the EM density map, and was therefore not included in the structure. **(c)** The model in *b*, rigid-body fitted into our map. The area in which Ac45 binds in the pre-existing V-ATPase structure (PDB: 9BRA) is indicated in the black box. **(d)** Comparison of the V-ATPase map from this study (centre) to the V-ATPase maps from purified synaptic vesicles in EMD-44855 (left) and EMD-18556 (right). The maps were rendered at thresholds which allowed their membranes to be visualised and compared

In order to establish whether the V-ATPase was complete, we generated a model of the *C. elegans* V-ATPase by fitting AlphaFold^19^ predictions of each reported subunit^20^ to the structure of *Mus musculus* V-ATPase (PDB: 9BRA) obtained in native membranes^21^. We then rigid-body fitted this model into our map (Figure 3c). All subunits in the V_1_ domain of the V-ATPase fit well into our map. However, we were not able to reliably fit the V_0_-associated accessory protein VHA-19 (Ac45) in the location observed in the *M. musculus* structure (Figure 3) (Supplementary Figure 5). AlphaFold predicts this subunit contains two domains which are connected by an extended flexible region. Such flexibility is expected to result in substantial positional variability between the domains, as shown by the large predicted alignment errors relative to one another in the AlphaFold prediction (>30 Å). Consistent with this, our map does not contain continuous or well-defined density corresponding to either the linker region or a fixed relative orientation of the two domains. This likely explains why the predicted structure cannot be fitted confidently into our reconstruction (Supplementary Figure 5). We also attempted to fit RAL-1, which is known to co-localise with AMSs and V-ATPases, into this region, but it did not reliably fit the density again due to the presence of predicted extended flexible regions. The resolution of the map in the membrane region did not allow us to confirm the precise stoichiometry of the c-ring; however, the arrangement of 9 VHA-1 (c) and 1 VHA-4 (c’’) subunits in our model fits the general shape of the density.

We performed 3D classification in order to investigate whether all particles were the complete, fully assembled V-ATPase. All classes were found to be the fully assembled V-ATPase, with both the cytosolic V_1_ and the transmembrane V_0_ domains present. We were unable to distinguish any molecules containing only V_0_ domains, which would represent the disassembled V-ATPase. As the surfaces of the AMSs are very crowded environments, it is possible that V_0_-only molecules are present in AMSs, but we could not effectively identify and pick them on the membrane surface.

We compared our *in situ* map of V-ATPase with other maps of V-ATPase obtained from purified, native vesicles from *M. musculus* (Figure 3d). All maps were structurally very similar, especially in the V_1_ domain, as expected given the high conservation of eukaryotic V-ATPases ^22^. However, one notable difference was the membrane curvature between our map and the other maps, which is unsurprising given the generally flat membrane geometry of the AMSs (Figure 3d).

### Distribution of V-ATPase on AMSs and their higher-order structure

A longstanding structural question concerning V-ATPases is whether they form higher-order oligomeric assemblies in native cellular environments. To date, no structural evidence has demonstrated such oligomerisation in V-ATPases, in contrast to the well-documented higher-order assemblies observed in ATP synthases^23–28^. Currently, other structures of V-ATPases have only shown monomers, but these structures have either been solved *in vitro* or in purified membranes, conditions which may have perturbed their native oligomeric state^29^. To address this, we analysed the spatial distribution of V-ATPases within AMSs to assess whether they adopt higher-order oligomeric states *in situ*.

Initially, we segmented the membrane of the AMSs in our tomograms and back-plotted the positions of each V-ATPase particle that contributed to the final map (Figure 4). V-ATPases are packed across the membrane surface of AMSs, with the exception of the most curved parts of the membrane folds (Figure 4a-b). Qualitatively, particles appear to cluster together, with some appearing to form dimers (Figure 4c). However, during the process of subtomogram averaging, we observed no evidence of dimerisation. Instead, we saw a ‘ring’ of density around the V-ATPase head at low thresholds (Figure 4d), indicating the presence of closely neighbouring particles located in random positions relative to the V-ATPase.

**Figure 4:**
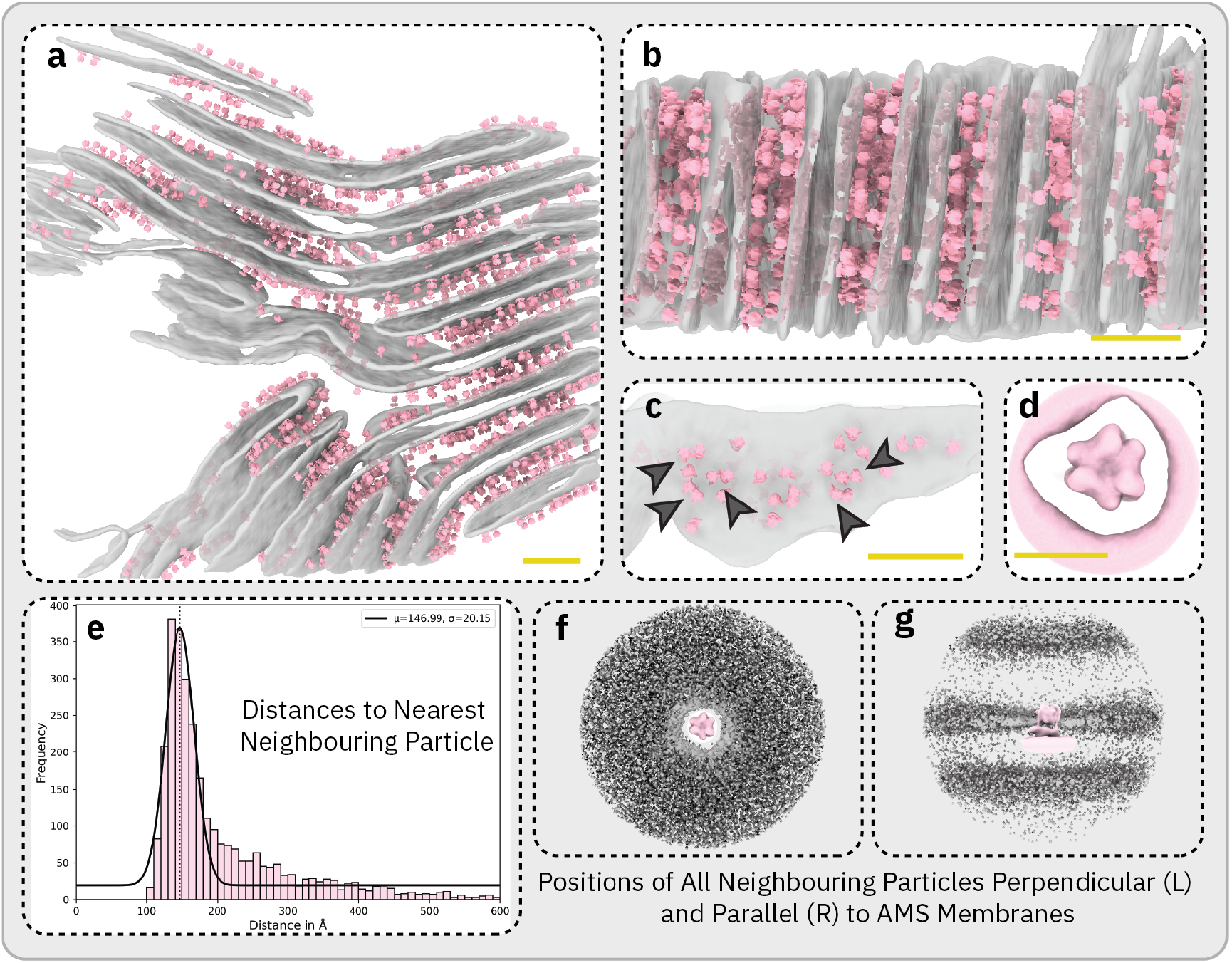
Analysis of V-ATPase distribution on AMSs. **(a)** Segmentation of a tomogram containing AMSs. Grey density corresponds to the AMSs membranes, and the pink particles correspond to V-ATPase particles which contributed to the final density map. Scale bar = 100 nm **(b)** The segmentation from *a* from a different angle. Scale bar = 100 nm **(c)** Close up of a single AMS membrane, with grey arrows pointing to V-ATPase dimer-like clusters. Scale bar = 100 nm. **(d)** The V-ATPase density map rendered at a low threshold to illustrate density corresponding to neighbouring V-ATPase particles. Scale bar = 150 Å **(e)** Nearest neighbour Analysis of V-ATPase particle positions after subtomogram averaging. A Gaussian curve was fitted to the data. The curve was centred at 147 Å. It should be noted that duplicate particles were deleted using a per-particle distance cut off of 100 Å during the subtomogram averaging workflow. Nearest neighbour analysis was also performed on particle coordinates prior to duplicate deletion to ensure that only duplicates were deleted. **(f)** Plot showing the distribution of all neighbouring particles relative to each particle. Each grey dot corresponds to the position of one neighbour. This view visualises the positions of neighbours on the same membrane from each reference particle. **(g)** Shows the data in *f* from a different angle so the positions of particles above and below V-ATPase particles, normal to the membrane, can be visualised.

To investigate this phenomenon further, we measured the distances from each picked particle to its nearest neighbouring V-ATPase (Figure 4e). This revealed a clear pattern of particles which have a neighbouring V-ATPase molecule within approximately 140-150 Å, which matches the distance between the V-ATPase head and the ring of surrounding signal seen in the map (Figure 4d). We also plotted the positions of every neighbouring V-ATPase particle to each picked particle in 3D (Figure 4f-g). This revealed that the positions and orientations of V-ATPases on the same membrane, and on neighbouring membranes, displayed no clear pattern relative to one another. Together, these data show that the V-ATPases in AMSs are not forming higher-order oligomers but are clustering closely together, possibly due to the density of V-ATPase packing on the membrane. However, it should be noted that there is a strong possibility that not all true positive V-ATPase particles were picked, either because they were missed during particle picking or because they were discared during subtomogram averaging 3D classification due to conformational heterogeneity. Consequently, it is possible that V-ATPases form larger particle clusters than those we observe here.

## Discussion

### The functional role(s) of AMSs

The precise role and function of AMSs remain unclear. One study suggests they could play a role in cell signalling in response to stress and damage, similar to the role of eisosomes in yeast^8^. However, the only evidence for this function is currently the co-localisation of phosphatidylinositol (4,5) bisphosphate (PIP_2_) to the AMSs. Whilst PIP_2_ is involved in signalling in response to stress, it also has diverse other functions^30^, including the recruitment of the V_1_ domain of V-ATPase to the membrane-embedded V_0_ domain^31^. As our map shows the V_1_ domain of V-ATPase is recruited to the V_0_ domain, it is likely the enrichment of PIP_2_ at the AMSs is a result of the signalling process required to assemble full V-ATPase molecules. Another suggested role is that the AMSs function as a mechanical attachment site between the epidermis and the cuticle^8^. This hypothesis is based on the observation that meisosomes occur in regions lacking hemidesmosomes, which are known to mediate cuticle–epidermis attachment^1^. However, the experimental evidence provided for this is limited. To establish a connection between AMSs and cuticle attachment, a previous study mutated furrow collagens in the cuticle and observed that the cuticle detached from the epidermis^8^. However, the correlation between this phenotype and AMSs is unclear. It is more likely that the detachment of the cuticle is a result of disruption between the known^32^ interactions of the collagen and hemidesmosomes. Other studies found that the V_0_ domain of V-ATPase, along with the GTPase RAL-1, is critical in the secretion of exosomes from MVBs in the AMSs, and other locations in the plasma membrane^5,7^.

In this study, we resolved the AMSs at a resolution that enabled us to visualise individual V-ATPase molecules lining the membrane folds. Previous studies have shown that V-ATPases can exist in two primary forms: (1) The complete, assembled V-ATPase which contains both the transmembrane V_0_ domain and the cytosolic V_1_ domain, and (2) the disassembled V-ATPase which only contains the V_0_ domain^33^. V-ATPase is known to act as an ATP-dependent proton pump responsible for acidifying intra- and extracellular compartments only when fully assembled^33^. However, for certain signalling and secretion functions, only the V_0_ unit is required to be present. In this study, we were only able to resolve the presence of completely assembled V-ATPases. Given their orientation along the AMS membrane folds, the fully assembled V-ATPases are positioned to pump protons directly into the cuticle, acidifying the extracellular space adjacent to the collagen fibres. The widespread presence of complete V-ATPase complexes across the AMS surface strongly suggests that proton pumping and extracellular matrix acidification represent a primary physiological function of AMSs which represents a role not previously attributed to these structures. The close proximity of mitochondria to AMSs may reflect the high local ATP demand required to sustain V-ATPase activity.

V-ATPases are known to play a key role in acidifying intracellular compartments such as lysosomes, vacuoles, and endosomes, as well as extracellular spaces^34^. However, the purpose of acidifying the cuticle is unclear. Previous studies showed proton-pumping by V-ATPases is key in regulating osmolarity in the excretory canal. V-ATPases could play a similar role at the cuticle^7,47^. Another previous study showed that the position of the AMSs is altered in moulting larvae so that the membranes align between cuticle furrows^8^. This indicates that AMSs may be involved in the moulting and/or cuticle remodelling process. Moulting requires the degradation of collagen, which constitutes a major component of the cuticle. A number of previous studies have established that the acidic environments created by V-ATPases are required for the breakdown of collagen by the cathepsin family of proteins^35,36^. Cathepsin gene expression has been observed at the cuticle of moulting C. elegans^37,38^. Consequently, we speculate that V-ATPases may play a role in acidifying the cuticle to facilitate cathepsin-mediated collagen degradation. However, further work is required to establish this relationship.

Both assembled and disassembled forms of V-ATPase have been implicated in signalling and/or secretion pathways. One protein associated with the exosome secretion pathway and known to co-localise to the AMSs is RAL-1. However, in this study, we could not resolve any protein density that could be unambiguously attributed to RAL-1, most likely due to its small size (29 kDa) that is incompatible with cryo-ET analyses. As a result, our structural data do not provide additional insights into the role of AMSs in exosome secretion. Future work could identify the precise location and distribution of RAL-1 using still-emerging protein tagging methods for correlative electron and light microscopy such as GEMs^39^ or ExoSloNano^40^. It has been previously reported that microtubules bind directly to the C subunit in the V_1_ domain of V-ATPases in yeast^41^. We also observed microtubules interacting with V-ATPases, confirming these interations also occur in *C. elegans*.

### The *in situ* structure of V-ATPase

The structural characterisation of membrane protein complexes within their native environment remains a central challenge in cell biology. Our *in situ* map of the *C. elegans* V-ATPase represents an important step in the structural understanding of this highly conserved proton pump, previously studied exclusively through *in vitro* systems ^21,42–50^. By visualising the fully assembled V-ATPase complexes within intact tissue, we demonstrate that V-ATPases are structurally conserved. Importantly, our data provide no evidence for the supramolecular assembly of V-ATPase in native conditions. Instead, our data show that V-ATPase particles often cluster on the AMSs, perhaps due to the fact that they are densely packed, but do not form oligomers with defined orientations relative to one another. These clusters had previously been observed in low abundance in purified synaptic vesicles^42^.

The lack of higher-order oligomers of V-ATPase is in direct contrast to the closely related mitochondrial ATP synthase. The dimerisation of ATP synthase plays a crucial role in imposing the shape of the mitochondrial cristae, as the dimer rows impose strong local curvature on the mitochondrial membrane^25,51^. This curvature drives the formation of sharply curved cristae ridges and the adjacent narrow cristae compartments. Such geometry is speculated to increase the efficiency of ATP production by concentrating protons in the intermembrane space which generates a larger proton gradient across the inner mitochondrial membrane^25^. The gap between the cuticle-facing folds of the AMS (∼30 nm) also forms a narrow, small region in which protons can accumulate. We speculate that V-ATPase proton pumping activity into this small space may make a highly-localised acidic environment within the cuticle, possibly to assist collagen degradation. In contrast to the mitochondrial membrane, this region is not created by membrane curvature induced by ATPases as the monomeric V-ATPases on AMSs exhibit no membrane bending properties and no V-ATPase molecules were observed at the inflexion point of the membrane folds. In order to determine the mechanisms involved in membrane folding during AMS production, future studies will need to ascertain the point in *C. elegans* development in which they are formed.

The absence of VHA-19 (Ac45) from our *in situ* V-ATPase map, despite its presumed presence in the fully assembled complex, underscores the limitations of sub-tomogram averaging in resolving conformationally heterogeneous or flexibly tethered domains. This reinforces the need for complementary approaches, such as *in situ* crosslinking and native mass spectrometry, to fully map the subunit composition and interaction networks of protein complexes such as V-ATPase. Similarly, the inability to localise RAL-1, a small GTPase implicated in V-ATPase-related trafficking pathways, reflects limitations of cryo-ET when working with small (<50 kDa) proteins.

Taken together, our study establishes a new workflow for understanding V-ATPase structure and organisation in a physiologically relevant setting. It opens avenues for future studies aimed at understanding the membrane re-modelling mechanisms leading to AMSs morphogenesis and how membrane architecture and protein clustering converge to support compartmentalised acidification in multicellular organisms.

## Acknowledgements

We thank members of the Mattei lab and the cryo-EM community at EMBL for useful discussions and technical support, in particular Zhengyi Yang and Georg Wolff. We would like to thank the Zimmer lab for contributing the *C. elegans* strain used in this study. We would also like to thank Fergus Tollervey and Yannick Schwab for critical readings of the manuscript, Rasmus Kjeldsen Jensen for providing the nearest neighbour analysis script, and Linhua Tai for providing a script to assist removing suboptimal images from tilt series within the WarpTools pipeline. EP was funded by the Deutsche Forschungsgemeinschaft (DFG, German Research Foundation) – SPP2416 project number 525894472, and the EMBL Interdisciplinary Post-doc Program (EIPOD), AB was funded by the European Union (GA#101094250), HVDR was supported by the EMBL International PhD program, and SM was funded by the European Molecular Biology Laboratory. We acknowledge the access and services provided by the Imaging Centre at the European Molecular Biology Laboratory (EMBL IC), generously supported by the Boehringer Ingelheim Foundation. This paper was typeset with the bioRxiv word template by @Chrelli: www.github.com/chrelli/bio-Rxiv-word-template

## Author contributions

Conceptualisation, EP and SM; sample preparation and serial lift-out, EP, AMS, and AB; cryo-ET data collection, EP; cryo-ET and STA data processing, EP; tomogram segmentation analysis, EP and HVDR; manuscript writing, EP; manuscript revisions, EP and SM; funding acquisition, SM.

## Data availability

The subtomogram averaging maps of the consensus V-ATPase (EMD-XXXXX), and the consensus ribosome structure (EMD-XXXXX) have been deposited in the Electron Microscopy Data Bank (EMDB). The atomic co-ordinates of the rigid-body fitted *C. elegans* V-ATPase model (PDB: XXXX) has been deposited in the Protein Data Bank (PDB).

## Materials and Methods

### *C. elegans* culture and high-pressure freezing

*C. elegans* expressing CHS-2::mMaple were cultivated according to standard methods on NGM^52^. We transferred 10 L4 worms to a fresh NGM plate 72 hours before high-pressure freezing. After 24 hours, the adult worms were removed. This resulted in a culture of L4-stage and early adult worms on the day of high-pressure freezing. In preparation for high-pressure freezing, we prepared an *E. coli* paste which was used as a cryo-protectant. To make this paste, we grew an overnight culture of *Escherichia coli* in 100 mL OP50 medium. We pelleted the cells by centrifugation at 5000 *xg* for 10 minutes. The pellet was resuspended in 450 μL of M9 buffer supplemented with 20% bovine serum albumin. The *E. coli* suspension was pelleted again by centrifugation at 5000 *xg* for 10 minutes. The pellet was resuspended in 50 μL of M9 buffer supplemented with 20% bovine serum albumin. To high-pressure freeze *C. elegans*, up to 50 worms were collected with a worm pick under a stereomicroscope and were added to the 100 μm well of a 3 mm Type A sample carrier (Leica Microsystems) together with 1.5 μL of *E. coli* paste. The engraved side of a TopFinder Type B carrier (Wohlwend) was used to seal the specimen before high-pressure freezing in a Leica EM ICE (Leica Microsystems).

### Fluorescence microscopy of high-pressure frozen carriers

Fluorescence microscopy of high-pressure frozen carriers was performed using a Zeiss LSM 900 Airyscan 2 microscope equipped with a cryo-stage (Linkam). Images were acquired using a 5x objective (0.4 NA) in widefield mode. A 488-nm laser was used to acquire images of CHS-2::mMaple fluorescence. Reflected-light images were also acquired to generate images of the surface of the ice. Illumination and detection were controlled by ZEN Blue software. Fluorescence and reflection images were superimposed in FIJI^53^.

### Serial Lift-Out of *C. elegans*

Serial lift-out was performed similar to a previously described workflow^10^. A high-pressure frozen carrier containing *C. elegans* worms that were deemed suitable for lift-out after fluorescence milling was identified. This carrier was loaded into a 35° pre-tilted Aquilos shuttle (Thermo Fisher Scientific) in cryo-conditions. A clipped copper rectangular pattern 400×100 mesh support grid (Agar Scientific) was loaded into the other shuttle slot for use as a receiver grid. The shuttle was loaded into an Aquilos 2 focused ion beam-scanning electron microscope (FIB-SEM) with an integrated fluorescence microscope and an EasyLift needle system. A copper block was attached to the needle from the receiver grid via copper redeposition. The worm identified for lift-out during previous fluorescence microscopy was located within the high-pressure frozen carrier using the integrated fluorescence microscope in the FIB-SEM. The sample was then protected from beam damage by applying a metal-organic platinum layer using the gas injection system (GIS). Trenches were milled surrounding the worm of interest using a current of 16 nA. An undercut of the block of ice containing the worm was then performed using a current of 3 nA. The copper block on the needle was attached to the block of ice containing the worm using copper redeposition. The block of ice was freed from the carrier by milling the last remaining attachment point of the block to the carrier. The needle was used to move the block of ice containing the worm to the receiver grid. The block was repeatedly milled using the FIB to deposit a series of 5 μm slices of worm containing ice onto the receiver grid with a current of 1 nA. These slices were attached to the receiver grid using copper redeposition on the sides of the slice as they were lowered into the gaps between the copper bars. Slices were imaged using the integrated fluorescence microscope to visualise the precise location of the worm within. Another protective metal-organic platinum layer was applied using the GIS. Slices were then manually thinned using decreasing milling currents depending on sample thickness. The following currents were used for each respective sample thickness: 1 nA to 1.5 μm, 0.5 nA to 1 μm, 0.3 nA to 0.7 μm, 0.1 nA to 0.5 μm, 50 pA to 250 nm. The resulting lamellae were polished to approximately 200 nm using 30 pA. Lamellae were then over-tilted by 0.5 ° and the back of the lamellae were further polished to achieve a more even thickness distribution across the ice. The lamellae were sputter coated at 10 mA for 3 seconds to reduce charging during cryo-ET^54^.

### Cryo-Electron Tomography Data Collection

The receiver grid from serial lift-out was loaded onto a Titan Krios G4 operated at 300 kV equipped with a Selectris X energy filter and a Falcon 4i direct electron detector (Thermo Fisher Scientific). Tilt series were acquired using SerialEM v4.1.15^55^ at a pixel size of 2.39 Å in nanoprobe mode with a 50 μm C2 aperture and a 70 μm objective aperture inserted. The energy filter slit was inserted at a width of 10 eV. Tilt series were acquired using a dose-symmetric tilt scheme^56^ with a 3° step with a total range of −51 to +51°, relative to the lamella tilt. The electron dose per tilt of either ∼3.93, ∼4.29, ∼4.48 e-/Å^2^, leading to a total tilt series electron dose of either ∼138, ∼150, ∼157 e-/Å^2^. A total of 280 tilt series were collected with PACE-tomo^57^.

### Cryo-Electron Tomography Data Processing

Tilt series were processed using the WARP/M pipeline for Linux^15,16^ with all steps being carried out using this software, unless stated otherwise. Initially,motion correction and CTF estimation were carried out. Tilt series stacks were then generated. These stacks were manually inspected and poor-quality images were removed from further processing. AreTomo2^58^ was used for initial tilt series alignment. AreTomo2 was used to generate preliminary tomograms which were used to estimate the thickness of each tomogram. These thickness values were used to set the appropriate-AlignZ value for each tilt series during another round of AreTomo2 tilt series alignment. The defocus hand was then estimated, and the tomograms flipped to ensure correct tomogram defocus handedness. 3D-CTF corrected tomograms were then generated at a pixel size of 14.34 Å/px. CryoCARE was used to denoise these tomograms before another round of denoising in IsoNet^59,60^. Denoised tomograms were solely used for manual visualisation and were not used in any data processing steps. A total of 216 tomograms were of a sufficient quality to analyse.

### Ribosome Subtomogram Averaging and Tomogram Optimisation

Ribosome particles were picked using template matching with PyTOM^61,62^ on 19.12 Å/px, 3D-CTF corrected tomograms. Initially, a plant ribosome was used as a template (EMD-15806)^63^ to pick on a subset of tomograms. The ribosomes picked from these tomograms were extracted as 3D particles by WARP^15^ and averaged in RELION (v4.0.1)^18^ to a resolution of ∼40 Å. This structure was used as a template in another round of PyTOM template matching on all tomograms. Ribosomes were also picked in another dataset from an additional lift-out session on the same high-pressure frozen carrier. Junk particles were removed using 2D and 3D classification before 3D auto-refinement which was repeated at progressively smaller pixel sizes until the resolution stopped improving. RELION v5.0.0^17^ was used for unbinned 3D auto-refinement due to the improved speed of refinement. Particles were then refined in M^16^ until the map was improved to a 5.6 Å resolution from a total of 72,793 particles. In M, only ctf_defocus, refine_particles, and refine_imagewarp 1×1 were used as refinement parameters. These 3 parameters optimise tomogram CTF estimation, ribosome particle poses, and tomogram tilt series alignment, respectively. 3D-CTF corrected tomograms were re-generated using the improved tomogram CTF estimation and tilt series alignment metadata from M. Tomograms were then denoised as before.

### V-ATPase Subtomogram Averaging

V-ATPase particles were manually picked using Surforama^64^ in combination with MemBrain-V2^65^. Particle coordinates were converted from .star files to Dynamo tables using relion2dynamo (https://github.com/EuanPyle/dynamo2relion). Particles were extracted from 3D-CTF corrected tomograms using Dynamo^66^. An initial model was generated by averaging all particles together using the Euler angles assigned during manual picking, in which the particles were oriented normal to the membrane. Particles were initially aligned to the membrane by masking the membrane in the initial model and only allowing particle translations in the direction of the membrane. Particles were then aligned using a limited cone range (5°) and limited translation shifts (2 px). The resulting table from this alignment was then converted from a Dynamo table to .star file using dynamo2relion (https://github.com/EuanPyle/dynamo2relion). WARP was used to extract 3D particles at 19.12 Å/px before alignment in RELION (v4.0.1)^18^. A custom script by Alister Burt was used to define the centre of the V-ATPase within the reference and to re-extract particles with V-ATPase centred within the box (https://gist.github.com/alister-burt/8744accf3f4696dd6d83fc9c4690612c). Particles were then extracted by WARP at a pixel size of 9.56 Å/px. In RELION, 3D classification with alignment and 3D auto-refinement was carried out to isolate a class of particles in which the V-ATPase peripheral stalks became visible. The resulting map was then used as a template for template matching in PyTOM^61^ on all AMS containing tomograms. To ensure every particle was picked, the cross-correlation threshold for determining whether a particle was picked or not was manually lowered to ‘over pick’ particles. Particles were extracted by WARP at a pixel size of 9.56 Å/px. A series of 3D classifications were carried out to remove false positive particles from template matching. One class from 3D classification was unambiguously V-ATPase with 3 clear peripheral stalks. This class was used for 3D auto-refinement until the resolution stopped improving. Particles were then refined in M^16^ only using refine_particles, and refine_imagewarp 1×1 as refinement parameters, as before in ribosome subtomogram averaging. Improved tomograms were reconstructed and the V-ATPase map from M was used as a template for another round of PyTOM template matching. Picks were processed as before and refined in RELION until resolution stopped improving. Particles were refined in M as before, except using refine_imagewarp 4×4, and then re-extracted for further refinement in RELION. A resolution of 17.4 Å was achieved from 2,988 particles.

### V-ATPase Rigid-Body Fitting

The subunits comprising the *C. elegans* V-ATPase were identified from a previous study^20^. The structure of each V-ATPase subunit was modelled using AlphaFold^19^. A series of V-ATPase models were docked into our V-ATPase map using *fitmap* in ChimeraX^67,68^. The model which fit our map best was the state 2 V-ATPase model from *M. musculus* (PDB: 9BRA)^21^. Each AlphaFold-predicted V-ATPase subunit from *C. elegans* was then aligned onto the *M. musculus* model using *matchmaker* in ChimeraX. This generated a model of the complete V-ATPase which was then docked into our V-ATPase map. Some regions of the model were deleted due to steric clashes. All subunits fit into our V-ATPase map well with the exception of VHA-19 which was consequently deleted from the model.

### Tomogram Segmentation and Analysis

IsoNet denoised tomograms were segmented using a combination of Mem-Brain and ArtiaX^65,69^. MemBrain was used to segment the membranes, collagen fibres, and microtubules within the tomograms. The final positions and orientations of ribosomes and V-ATPases were read from the .star files produced at the end of their respective subtomogram averaging processing and back-plotted into the segmentation using ArtiaX. Segmentations were visualised within ChimeraX^67,68^. To calculate the distances between AMS membranes, we imported the MemBrain segmentations of the AMS membranes into DragonFly^70^. Segmentations of membranes which were not continuous, or otherwise incomplete, were excluded from further analysis. Membranes belonging to the same fold were grouped together as connected components. The largest component in each tomogram was used as a reference from which the distance of all other components was measured from. This analysis was performed individually for each tomogram.

**Supplementary Figure 1:**
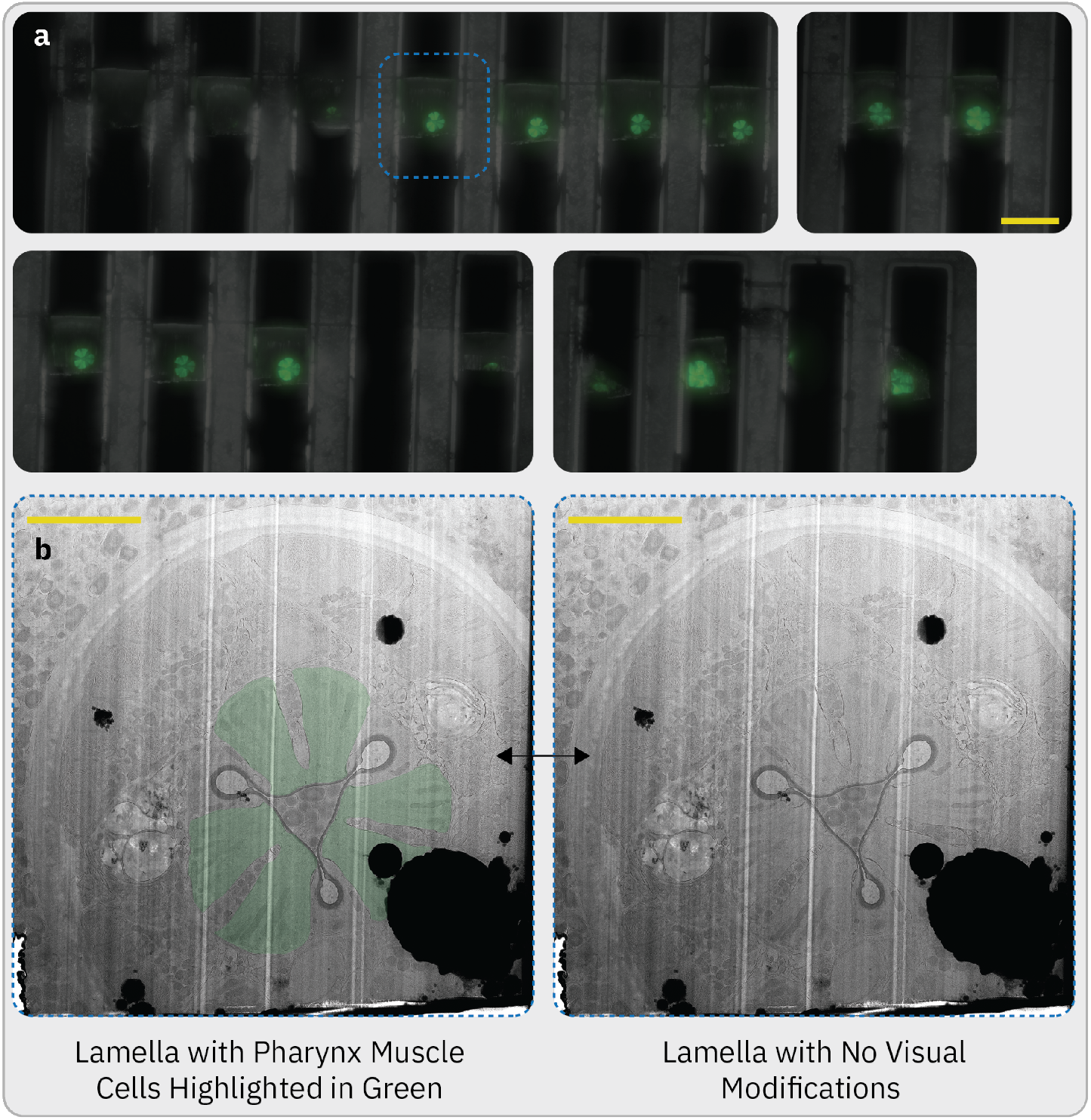
CHS-2 is localised within pharynx muscle cells. **(a)** Widefield microscopy image of deposited lamella with overlaid cryo-fluorescence signal of CHS-2::mMaple inside the FIB-SEM. Whilst there are 16 lamellae present here, only 11 were of sufficient quality for imaging by cryo-ET due to ice contamination or due to lamella breakages during sample transfer to the TEM. The lamella visualised in *b* is highlighted in the blue box. Scale bar = 40 μm. **(b)** TEM overview images of the lamella indicated in *a*, with (left) and without (right) the pharynx muscle cells highlighted in green. The shape and location of the mMaple fluorescence in *a* match the location of the pharynx muscle cells in *b*. Scale bar = 5 μm.

**Supplementary Figure 2:**
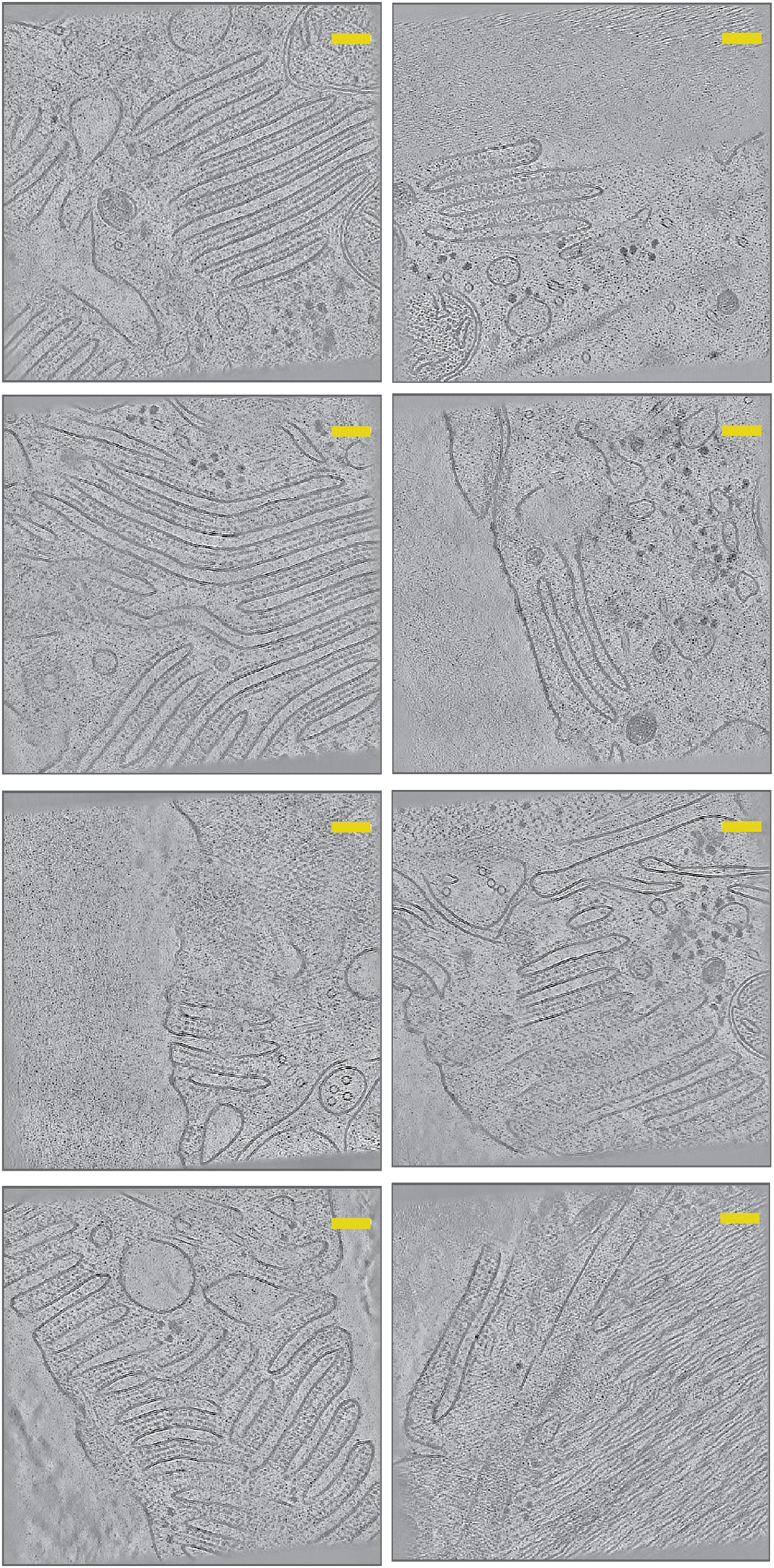
Montage of all tomograms collected that contain AMSs. Single slices of each tomogram in which AMSs were positively identified. Scale bar = 100 nm.

**Supplementary Figure 3:**
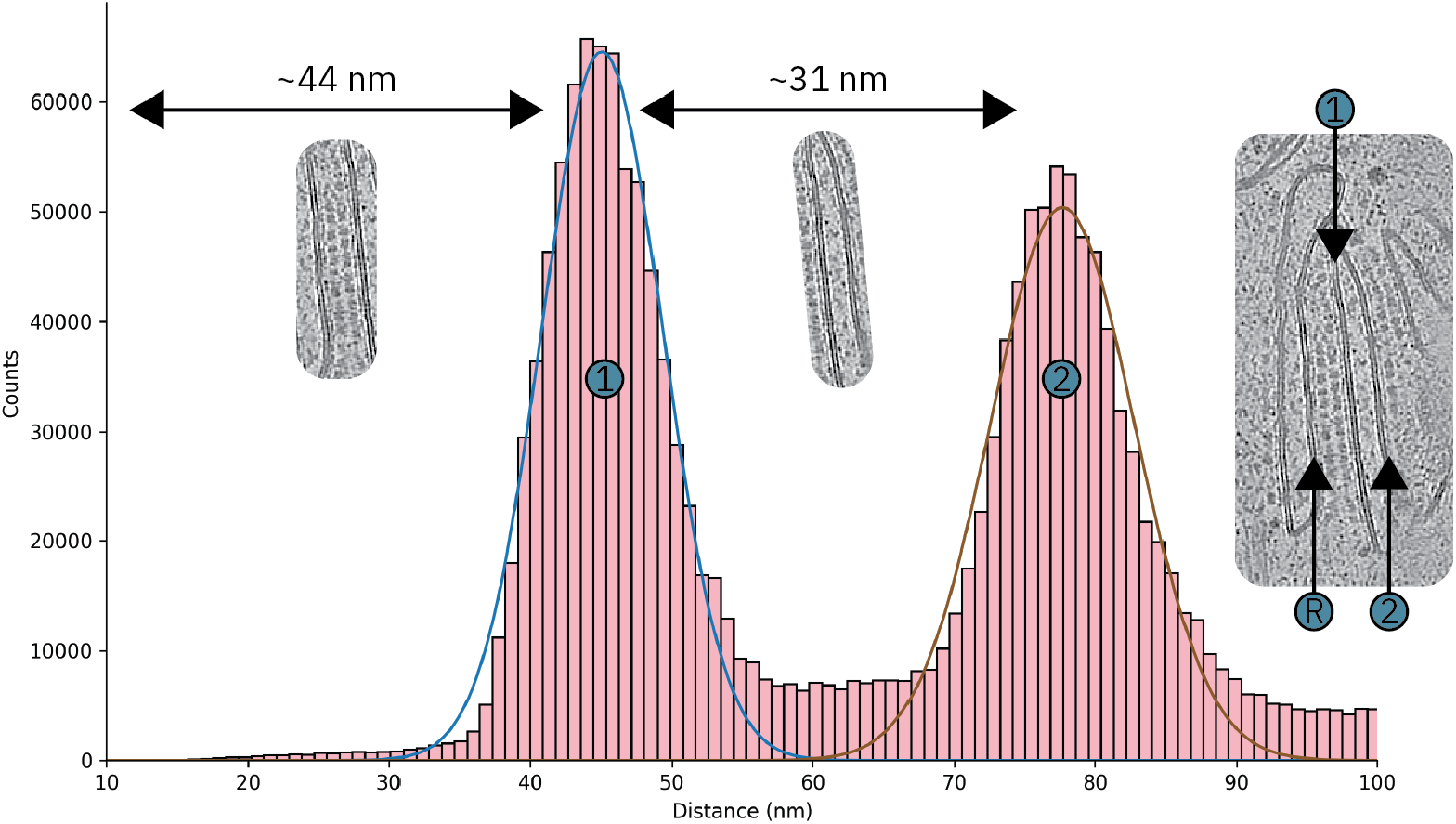
Dimensions of the AMSs. Histogram from one tomogram measuring distances between AMS membranes. (R) designates an example of a reference membrane, from which the distances of membranes (1) and (2) are measured. Cytoplasmic facing membranes and cuticle facing membranes were calculated to be 44 nm and 31 nm apart, respectively, averaged across all tomograms.

**Supplementary Figure 4:**
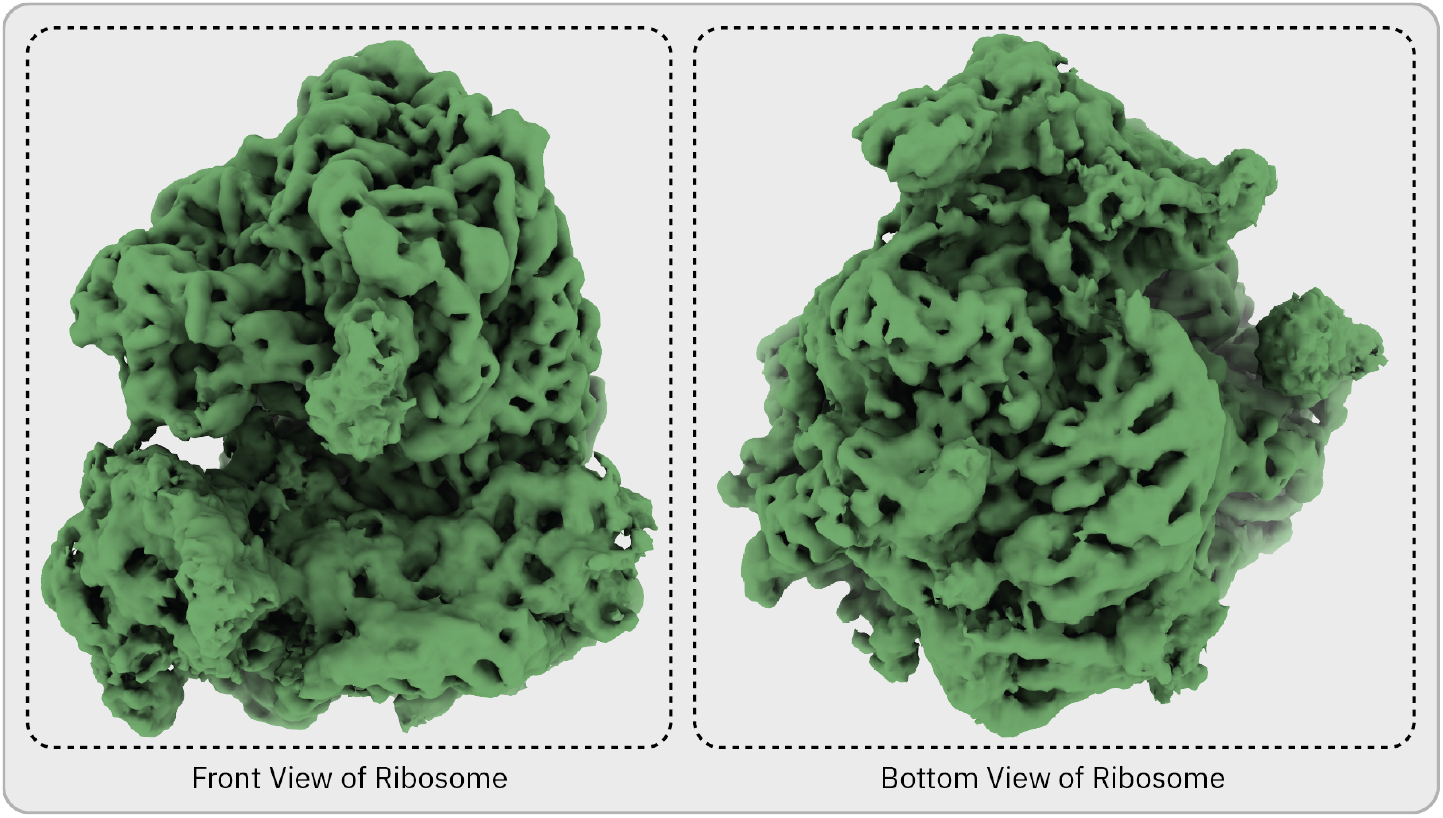
Structure of the ribosome from *C. elegans*. A 5.6 Å density map of a ribosome obtained *in situ* from *C. elegans* tomograms viewed from two different angles. This map was refined in M^15^ using an image warping parameter of 1×1, alongside particle and CTF refinement, in order to improve the quality of the tomograms. Ribosome particles were averaged from tomograms both with and without AMSs.

**Supplementary Figure 5:**
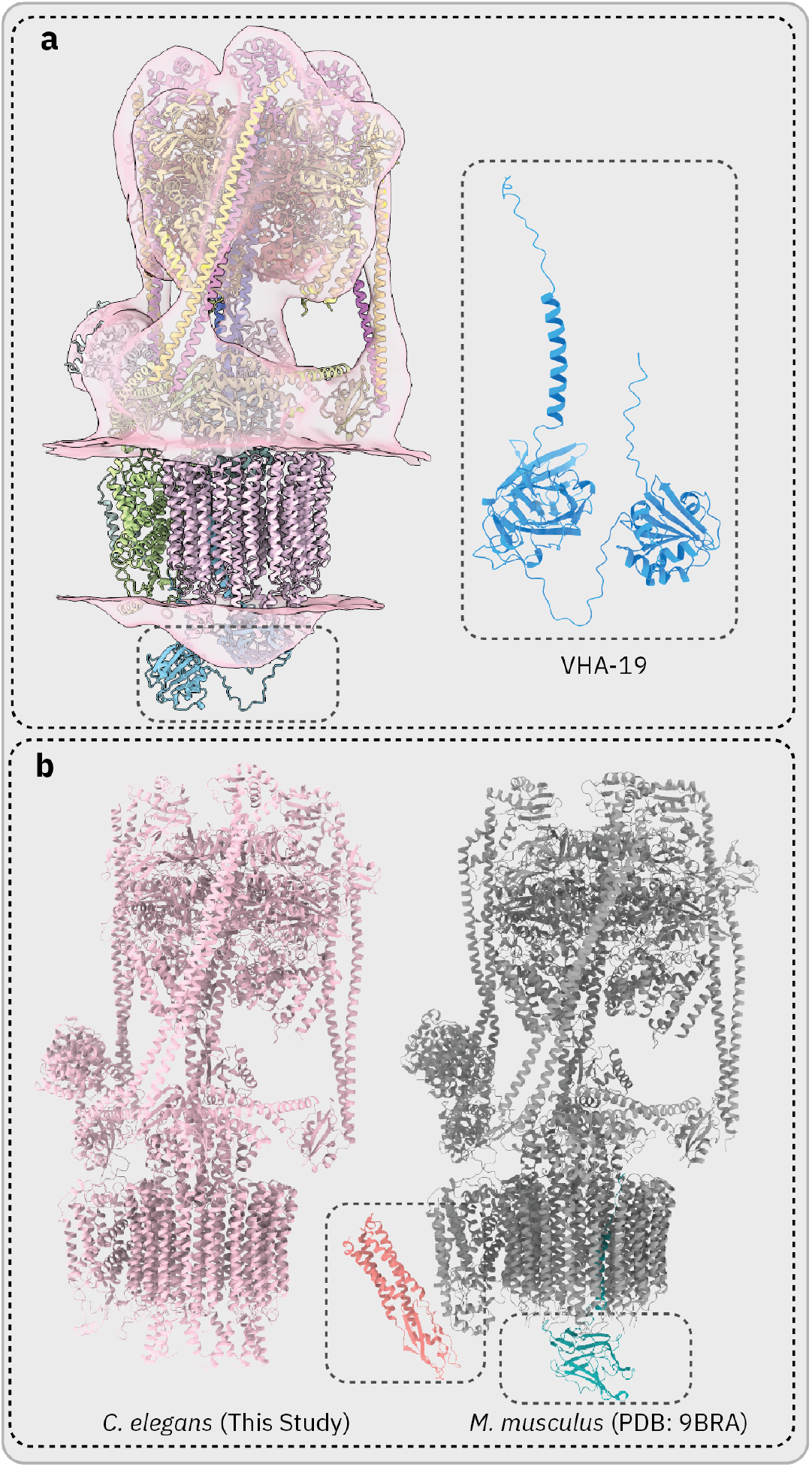
Analysis of *C. elegans* V-ATPase Model. **(a)** Rigid body fitting of VHA-19 into the V-ATPase map. The black box (bottom left) highlights the area in which VHA-19 is proposed to bind. The inset shows a close up of the AlphaFold predicted VHA-19 structure. Two domains separated by a flexible region are clearly visible, alongside a transmembrane helix. **(b)** Comparison of the models of V-ATPase in *C. elegans* (left) and *M. musculus* (right, PDB: 9BRA). Two subunits are present in the *M. musculus* model which are not present in the *C. elegans* model. These subunits are Ac45 (VHA-19 homolog) (green) and synaptophysin (orange). Synaptophysin is a protein found in synaptic vesicles; therefore, it is not expected to be present in V-ATPases in C. elegans AMSs.

